# Context-dependent NMDA receptor dysfunction predicts seizure treatment in mice with human GluN1 variant

**DOI:** 10.1101/2024.11.01.619946

**Authors:** Sridevi Venkatesan, Daria Nazarkina, Megan T Sullivan, Yao-Fang Tan, Sarah Qu, Amy J. Ramsey, Evelyn K. Lambe

## Abstract

Mutations in N-Methyl D-Aspartate receptors (NMDARs) cause epilepsy and profound cognitive impairment, though the underlying subunit-specific vulnerabilities remain unclear. We investigate the impact of a severe human variant in the lurcher motif of obligate GluN1 NMDAR subunit using transgenic mice, leveraging context-specific dysfunction to devise a surprising treatment. We show that the GluN1 Y647S variant significantly reduces current flow through isolated NMDARs in the mouse brain. However, this loss-of-function paradoxically extends NMDAR-dependent dendritic integration, causing prolonged circuit-wide excitation that promotes seizures. Mutant receptors fail to sufficiently engage opposing dendritic ion channels that normally prevent NMDAR overactivation. Boosting negative feedback restores normal dendritic integration and successfully treats seizures *in vivo,* despite loss-of-function of isolated NMDARs. We demonstrate how seizures arise from loss-of-function NMDARs and target the interaction between a GluN1 variant’s receptor-level effects and its dendritic environment to treat them effectively.

## Introduction

N-Methyl-D-Aspartate receptors (NMDARs) are heterotetrametric ligand gated ion channels activated by the excitatory neurotransmitter glutamate^1,2^. NMDARs are vital for normal brain development and physiology and their disruption is implicated in multiple neurobiological disorders^3–5^. *De novo* missense mutations in the *GRIN* family of genes encoding for different NMDAR subunits cause a rare neurodevelopmental disorder characterized by epilepsy, developmental delay and intellectual disability^6–8^.

NMDAR patient variants have been previously examined using heterologous expression systems including oocytes^9–12^, HEK cells^13–15^, or viral expression in brain slice cultures^16^. These approaches characterize mutant NMDARs in artificial environments, classifying variants as loss-or gain-of-function based on the ion channel properties. It is increasingly becoming evident that comprehensive assessment in intact neural circuits using transgenic mice is further necessary to understand and treat the neurological consequences of NMDAR variants^15–19^. Particularly, human variants in the obligate GluN1 NMDAR subunit that cause severe neurological deficits in patients^8,20^ are yet to be examined.

NMDAR function is typically assessed at the level of isolated synaptic receptors^15,16,21^. However, dendritic NMDARs acting in an integrative capacity are increasingly recognized as critical for healthy brain function and cognition^22–26^. Critically, disruptions to NMDARs can result in heterogenous deficits across sub-cellular compartments^27,28^. Therefore, an important next step is to examine context-specific effects of NMDAR variants to determine effective treatments.

We investigate context-specific NMDAR dysfunction caused by a human variant in the obligate GluN1 NMDAR subunit using transgenic mice with the Y647S mutation^29^. This mutation occurs in a highly conserved transmembrane region of the NMDA receptor known as the lurcher motif^30^. Heterologous expression studies report reduced surface expression of mutant NMDARs, but variable changes in channel properties, and are unclear on whether this results in loss or gain-of-function of the receptor^6,20,29,31,32^.

We show that *Grin1* Y647S^+/-^ mice have reduced isolated NMDAR currents yet display paradoxically enhanced NMDAR-dependent integration that causes seizure-like events in brain slices. Compromised current through mutant NMDARs fails to engage negative feedback via calcium-activated potassium channels that typically prevent NMDAR overdrive^33^. We mimic restoration of negative feedback by raising brain magnesium levels and show that chronic oral treatment with magnesium L-threonate (MgT)^34^ significantly ameliorates seizure occurrence and severity in Y647S^+/-^ mice.

Our work reveals an unexpected mechanism for epilepsy caused by a loss-of-function *GRIN1* variant and highlights dietary magnesium supplementation as an effective treatment. We identify critical differences in the impact of a genetic NMDAR variant at isolated and integrative levels, showing that functional context is essential to determine treatments.

## Materials and Methods

### Animals

Mice heterozygous for the Y647S mutation in the *Grin1* gene encoding for the obligate GluN1 subunit (*Grin1* Y647S^+/-^ mice) were generated as described previously^29^. Thy1-GCaMP6f mice (RRID: IMSR_JAX:02828) were crossed with *Grin1* Y647S^+/-^ mice and the offspring which were positive for Thy1-GCaMP6f and Y647S^+/-^ or WT were used for widefield calcium imaging and subsequent *in vivo* treatments. All experiments were approved by the Temerty Faculty of Medicine Animal Care and Use Committee and followed Canadian Council on Animal Care and AVMA guidelines. Mice were group-housed and kept on a 12-hour light cycle with food and water access *ad libitum*.

### Brain slicing

Cortical brain slices were obtained as described previously^28,35,36^. In brief, the brain was rapidly extracted in ice cold sucrose ACSF (in mM: 254 sucrose, 10 d-glucose, 26 NaHCO_3_, 2 CaCl_2_, 2 MgSO_4_, 3 KCl and 1.25 NaH_2_PO_4_) and slices (400 μm thick) allowed to recover for ∼ 2 hours in oxygenated (95% O_2_, 5% CO_2_) ACSF (in mM: 128 NaCl, 10 D-glucose, 26 NaHCO_3_, 2 CaCl_2_, 2 MgSO_4_, 3 KCl, and 1.25 NaH_2_PO_4_) at 30°C. For whole-cell electrophysiology or wide-field calcium imaging, brain slices were transferred to the stage of a BX51WI microscope (Olympus, Tokyo, Japan) and perfused with oxygenated ACSF at 30°C. Recording electrodes (2–4 MΩ) filled with patch solution were used to record from layer 5 (L5) pyramidal neurons. Multiclamp 700B amplifier with Digidata 1440A and pClamp 10.7 software (Molecular devices) were used for data acquisition at 20 kHz. All recordings were compensated for liquid junction potential.

### Whole-cell electrophysiology to assess NMDAR function

Detailed experimental procedures including recipes for various patch solutions, protocols to evoke and measure excitatory postsynaptic currents, NMDAR plateau potentials, whole-cell NMDA currents, and NMDAR current-voltage curves are provided in Supplemental Methods.

### Widefield calcium imaging

To characterize the circuit-wide effect of the *Grin1* Y647S mutation, an optical measure of population level neuronal activity was recorded from WT and Y647S^+/-^mice expressing GCaMP6f under the *Thy1* promoter (*Thy1*-GCaMP6f) in layer 5 pyramidal neurons. Population neural activity in cortical brain slices was evoked by apical dendritic stimulation (10 pulses, 50 Hz) in layer 2/3 and measured using the Prime BSI Express camera (*Teledyne*) at 20 ms exposure through the 10X lens of the widefield microscope. Details of the analysis are provided in Supplemental Methods.

### Magnesium threonate treatment and seizure characterization

*Grin1* Y647S^+/-^ and WT littermate animals (84 ± 4 days old) were given either 0.5% w/v Magnesium L-Threonate (*AK scientific*) dissolved in their drinking water (MgT) or regular tap water (Control). Mice had access to water ad libitum and the MgT dose was calculated to be 1086 mg/kg (for 23g mice drinking 5ml per day). This equates to an elemental magnesium dose of 88.5 mg/kg per day. Mice were picked up and handled by the experimenter once a week at the same time of day (11 AM) for ∼ 2-3 minutes, followed by weighing the mice. Seizures occurring during this period were quantitatively assessed from video recordings of Y647S^+/-^ mice from both MgT and control groups by an independent observer. Mice underwent cage changes and water bottle switches on the day following the handling induced seizure assessment. After 12 weeks, control Y647S^+/-^ mice that displayed prominent seizures were switched to MgT treatment and their seizure incidence was monitored for the next 6 weeks. Videos containing visually identifiable seizures were assessed from weeks 6-12 of the treatment period. In total, 50 videos were analyzed and total seizure duration in seconds, exhibited symptoms, and seizure severity were measured as per our modified Racine scale^37,38^ (0 -normal behaviour, 1 - behavioural arrest and vacant stare, 2 - head nodding, facial automatisms, tail extension, 3 - unilateral clonus, 4 - bilateral clonus, rearing and falling, 5 - loss of posture). All animals were humanely euthanized after the experiment and re-genotyped.

### Open Field behaviour testing and tracking

On the 7^th^ week of MgT treatment, mice were handled for ∼ 2 minutes for seizure assessment and then placed in a circular open field arena where they were allowed to explore for 10 minutes under continuous overhead camera monitoring. The arena was cleaned thoroughly between each mouse. DeepLabCut^39^ and Bonsai^40^ were used to track animal body parts. Total distance traveled and velocity of the mice were calculated using DLCAnalyzer^41^.

### Statistics

Statistical tests were performed in Prism 9 (Graphpad). Data are presented as mean ± SEM. Comparisons of electrophysiological properties between WT and Y647S^+/-^ mice used two-tailed unpaired t-tests or Welch’s test if the variance was different between the groups. Two-way ANOVA or two-way repeated measure (RM) ANOVA with Sidak’s post hoc test were used when considering the interaction between genotype and other variables such as stimulus intensity or pharmacological blockers. Within-cell effects of pharmacological manipulations were evaluated using paired t-tests. One-way ANOVA with Tukey’s post hoc test was used to evaluate efficacy of treatments in restoring Y647S^+/-^ properties to WT levels. Distribution of running velocity across groups of mice was compared using Kruskal Wallis test. \**P* < 0.05, \*\**P* < 10^-2^, \*\*\**P* < 10^-3^, \*\*\**P* < 10^-4^

## Results

We investigate the neurophysiological consequences of the Y647S *GRIN1* patient variant that occurs in the transmembrane region of the GluN1 NMDA receptor (NMDAR) and causes intellectual disability and epilepsy^6,7,20^ in the patient. Using a mouse model (*Grin1* Y647S^+/-^ mice) that replicates the phenotype of epilepsy and cognitive deficits^29^, we examine prefrontal neurotransmission and isolated versus integrative NMDAR signaling to decipher and treat context-specific NMDAR dysfunction (**Fig 1A-C**).

**Figure 1.**
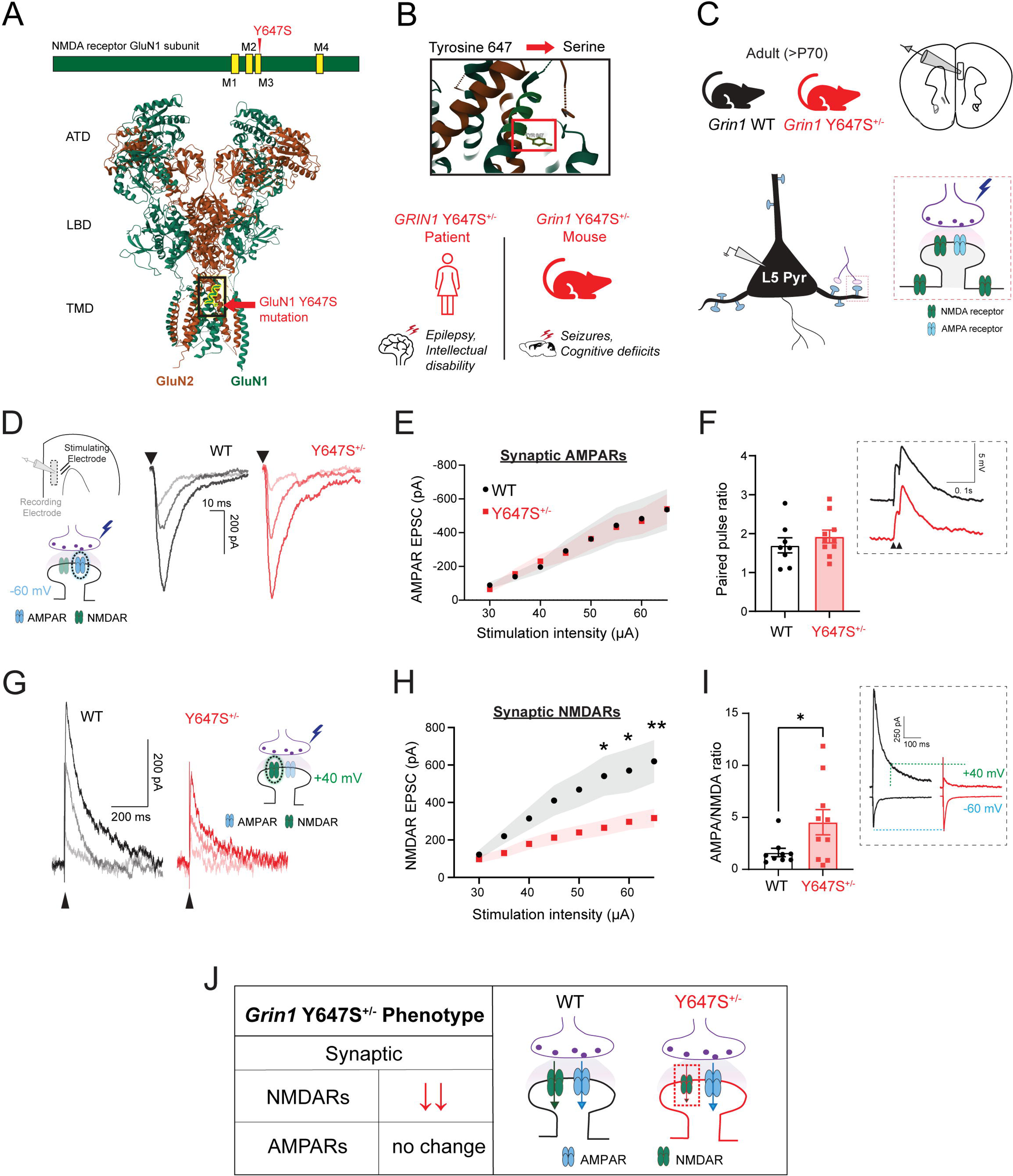
Isolated NMDA receptors show loss of function in mice with GluN1 Y647S^+/-^ patient variant. **A,** Schematic showing the position of the Y647S mutation in the M3 transmembrane domain of the GluN1 NMDA receptor (NMDAR) subunit (top), Structure of the human GluN1-GluN2A NMDAR tetramer (PDB accession number: 6IRF)^90^ highlighting the Y647 site in the channel pore (bottom). **B,** Tyrosine residue at the 647^th^ position of GluN1 is mutated to Serine, causing severe seizures and intellectual disability in the patient. Mutant mice (*Grin1* Y647S^+/-^) replicating the patient phenotype were used for slice electrophysiology. **C,** Coronal prefrontal cortex slices from adult wildtype (WT) and mutant (Y647S^+/-^) mice were used to record synaptic currents evoked by electrical stimulation of glutamate release in layer 5 pyramidal (L5 Pyr) neurons. **D,** Schematic with position of stimulating and recording electrodes in the brain slice. Isolated AMPAR excitatory postsynaptic currents (EPSCs) measured using cesium gluconate patch pipettes in the presence of GABAR blockers at -60 mV in WT and Y647S*^+/-^* at increasing stimulus strengths. **E**, AMPAR EPSC peak amplitude in WT and Y647S*^+/-^* at different stimuli are identical. **F,** Paired pulse ratio at 50 ms interval is similar in WT and Y647S*^+/-^* neurons. **G,** Isolated NMDAR EPSCs measured with AMPARs blocked at +40 mV in WT and Y647S*^+/-^*. **H**, NMDAR EPSC peak amplitude is reduced in Y647S^+/-^ neurons (\**P* < 0.05, \*\**P* < 0.01, Sidak’s post hoc). **I**, AMPA to NMDA ratio in WT and Y647S^+/-^ neurons (\**P* < 0.05, unpaired t-test). (Inset), EPSCs at -60 and +40 mV in WT and Y647S^+/-^, dotted line indicates NMDAR component at +40mV. **J,** Summary schematic showing preserved AMPAR but reduced NMDAR currents in *Grin1* Y647S^+/-^ mice. Bars denote mean ± SEM.

### Impact on prefrontal glutamate synapses

#### Intact AMPAR transmission in *Grin1* Y647S+/-mice

To determine the effect of the *Grin1* Y647S variant on glutamatergic synapses in intact neural circuits, we measured excitatory postsynaptic currents (EPSCs) in principal neurons in response to electrically evoked glutamate release. AMPA receptor (AMPAR) EPSC amplitude was similar between WT and Y647S^+/-^ (**Fig 1D-E**, F _(1, 143)_ = 0.001, *P* = 0.98, N= 7, 8 mice for WT and Y647S). Paired pulse ratio of postsynaptic potentials showed no difference between WT and Y647S^+/-^ neurons (**Fig 1F**, t _(16)_ = 0.90, *P* = 0.38). Average miniature EPSC amplitude and frequency measured to assess spontaneous glutamate release were also similar between WT and Y647S^+/-^ neurons (frequency: t _(12)_ = 1.21, *P* = 0.25, and amplitude: t _(12)_ = 0.07, *P* = 0.94). We conclude that presynaptic glutamate release probability and postsynaptic AMPAR strength are not altered by the Y647S variant.

#### Loss of function of isolated NMDA receptors in *Grin1* Y647S^+/-^ mice

Next, we directly assessed postsynaptic current through isolated NMDARs. NMDAR EPSCs were substantially reduced in Y647S^+/-^ neurons, with 50% reduction in peak amplitude at the highest tested stimulus (**Fig 1G-H**. 2-way ANOVA, genotype: F _(1, 192)_ = 37.78, *P* < 10^-4^; t_(192)_ = 3.11, \**P* = 0.017, Sidak’s post hoc). Reduction in Y647S^+/-^ NMDAR EPSC amplitude was observed in both sexes, with no significant effect of sex (F _(1, 22)_ = 0.005, *P* = 0.94). Y647S^+/-^ NMDAR EPSCs also showed faster decay kinetics with significant reduction in the half-width (WT: 94 ± 8 ms, Y647S^+/-^: 64 ± 8 ms, t _(16)_ = 2.64, *P* = 0.018). D-APV was equally effective at blocking NMDARs in both genotypes (2-way RM ANOVA, APV: F _(1, 10)_ = 49.99, *P* < 10^-4^) indicating similar receptor composition and antagonist efficacy.

To evaluate the relative loss of NMDAR function with respect to AMPARs within the same cell, we calculated AMPA/NMDA ratio (**Fig 1I**, inset). Consistent with the preservation of AMPAR but reduction in NMDAR responses, Y647S^+/-^ neurons showed significantly increased AMPA/NMDA ratio (WT: 1.6 ± 0.4, Y647S^+/-^ = 4.5 ± 1.2, t _(17)_ = 2.18, *P* = 0.04). In summary, the Y647S mutation in GluN1 subunits results in loss of isolated synaptic NMDAR currents in excitatory neurons without altering synaptic AMPARs or glutamate release probability (**Fig 1J**).

### Impact on integrative NMDAR signaling

#### Paradoxically prolonged dendritic excitation in *Grin1* Y647S^+/-^ mice

To determine the impact of the Y647S mutation on integrative NMDAR signaling, we examined NMDAR plateau potentials. Plateau potentials are NMDAR-dependent dendritic phenomena caused by glutamate spillover during repetitive stimulation and are vital for cognitive integration^42,43^ (**Fig 2A**). NMDAR plateau potentials elicited by 50Hz stimulation (10 pulses) showed a significant reduction in peak amplitude in Y647S^+/-^ neurons (**Fig 2B**, 2-way ANOVA, genotype: F _(1, 266)_ = 17.95, *P* < 10^-4^, N = 10,12 mice for WT and Y647S). This decrease in Y647S^+/-^ plateau amplitude was smaller (36% decrease, t _(266)_ =2.80, *P* = 0.038) compared to previously measured reduction in isolated synaptic NMDAR currents (50%).

**Figure 2.**
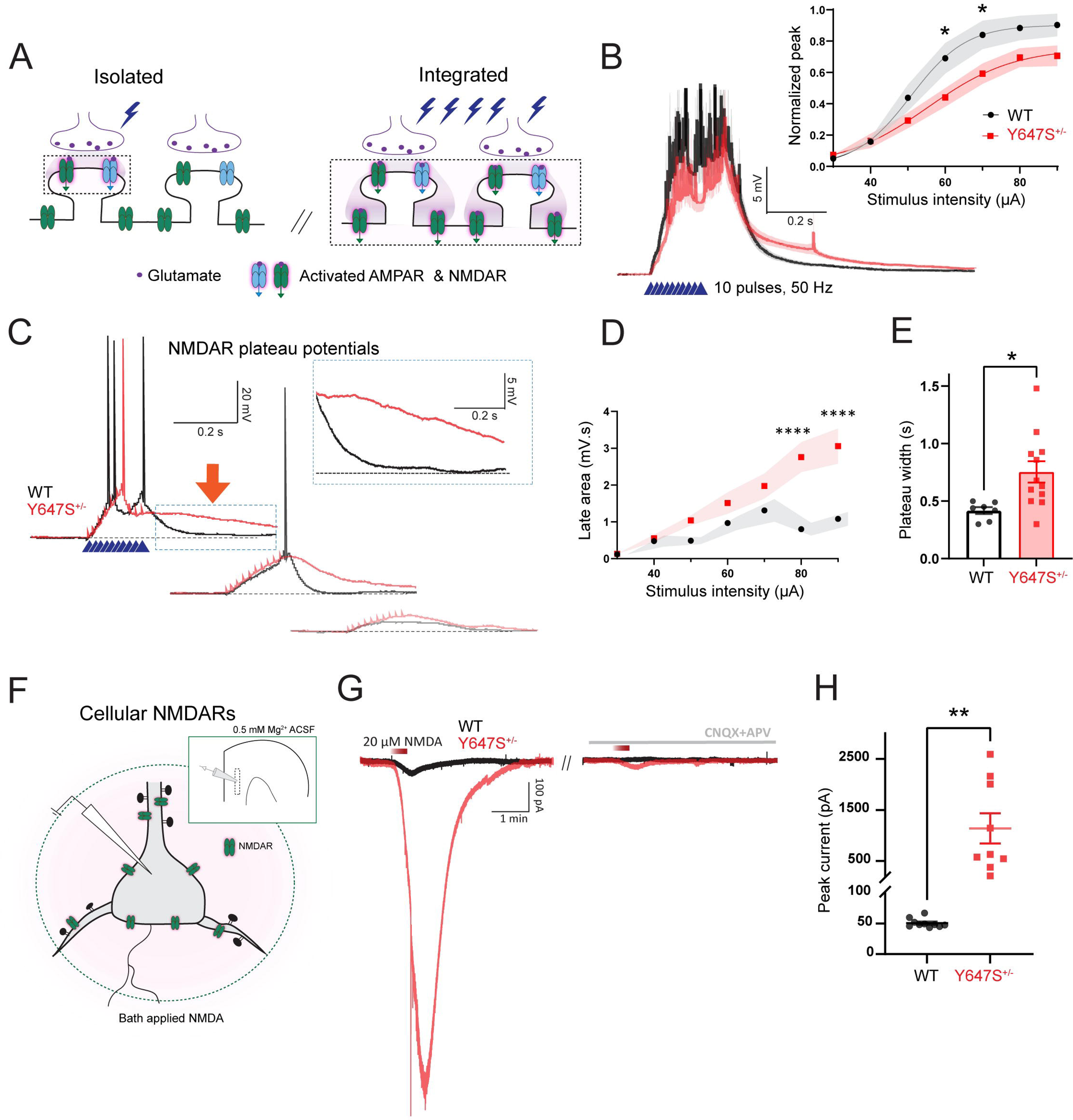
Paradoxically prolonged NMDAR integration and enhanced whole-cell NMDA currents in *Grin1* Y647S^+/-^ mice. **A,** Schematic showing that a single stimulus evokes limited glutamate release that activates only synaptic receptors, while repetitive high frequency stimulation causes glutamate spillover onto the dendrite, generating a dendritic plateau potential. **B**, Average NMDAR plateau potentials in WT and Y647S^+/-^ neurons evoked by 50 Hz stimulation, measured in the presence of AMPAR and GABAR blockers. (Inset) Normalized plateau potential amplitude at different stimulus intensities in WT and Y647S^+/-^ (\**P* < 0.05, Sidak’s post hoc). **C,** NMDAR plateau potentials show extended duration in Y647S^+/-^ neurons compared to WT at multiple stimulus intensities. **D,** Area under the plateau potential for the 1s period post-stimulation (late area) at different stimuli in WT and Y647S^+/-^ (\*\*\*\**P* < 10^-4^, Sidak’s post hoc). **E,** Total duration of the plateau potential (at 50 µA stimulation) in WT and Y647S^+/-^ neurons (\**P* < 0.05, unpaired t-test). **F,** Schematic shows the measurement of whole-cell NMDA currents in L5 neurons by the application of 20 µM NMDA. **G,** Whole-cell NMDA currents measured at -75 mV in WT and Y647S^+/-^ neurons in the presence of AMPAR blockers (left) are blocked by the NMDAR antagonist d-APV (right). **H**, Whole-cell NMDA current peak in WT and Y647S^+/-^(\*\**P* < 0.01, unpaired t-test). Bars denote mean ± SEM.

Despite their reduced amplitude, the duration of NMDAR plateau potentials was significantly prolonged in Y647S^+/-^ neurons, lasting over 200 ms beyond the WT neurons (**Fig 2C**). The area under the plateau potential for the 1s period post-stimulation (late area) was significantly greater (181% increase) in Y647S^+/-^ neurons (**Fig 2D**, 2-way ANOVA, genotype: F _(1, 127)_ = 36.74, *P* < 10^-4^; t_(127)_ = 5.33, *P* < 10^-4^, Sidak’s post hoc, N = 5, 7 mice for WT and Y647S). The total duration of plateau potentials was significantly increased in Y647S^+/-^ neurons (0.75 ± 0.09 s, **Fig 2E**) compared to WT (0.42 ± 0.03 s, t _(17)_ = 2.69, *P* = 0.015). NMDAR plateau potentials were blocked by the NMDAR antagonist d-APV (93% reduction) in both genotypes (2-way RM ANOVA, APV: F _(1, 8)_ = 35.21, *P* <10^-3^). The prolonged tail of Y647S^+/-^ plateau potentials was also sensitive to d-APV (72% reduction, t _(8)_ = 6.34, *P* = 0.0004, Sidak’s post hoc), indicating that it is an NMDAR-dependent phenomenon.

#### Unrestrained pharmacological NMDA currents in *Grin1* Y647S^+/-^ mice

Next, we measured the cellular impact of the Y647S mutation by recording whole-cell currents evoked by exogenous application of 20 µM NMDA (**Fig 2F**). Y647S^+/-^ neurons show greatly amplified NMDA currents (1138 ± 235 pA, unpaired t-test: t _(17)_ = 3.89, *P* = 0.001) that appear to be unrestrained compared to WT (51 ± 2 pA, **Fig 2G-H**). NMDA currents were blocked by d-APV in both genotypes (Fig 3G, 91% and 98% reduction in WT and Y647S^+/-^ respectively). Enhanced NMDA currents were observed in Y647S^+/-^ neurons of both sexes, with no significant effect of sex (F _(1, 15)_ = 1.307, *P* = 0.27).

**Figure 3.**
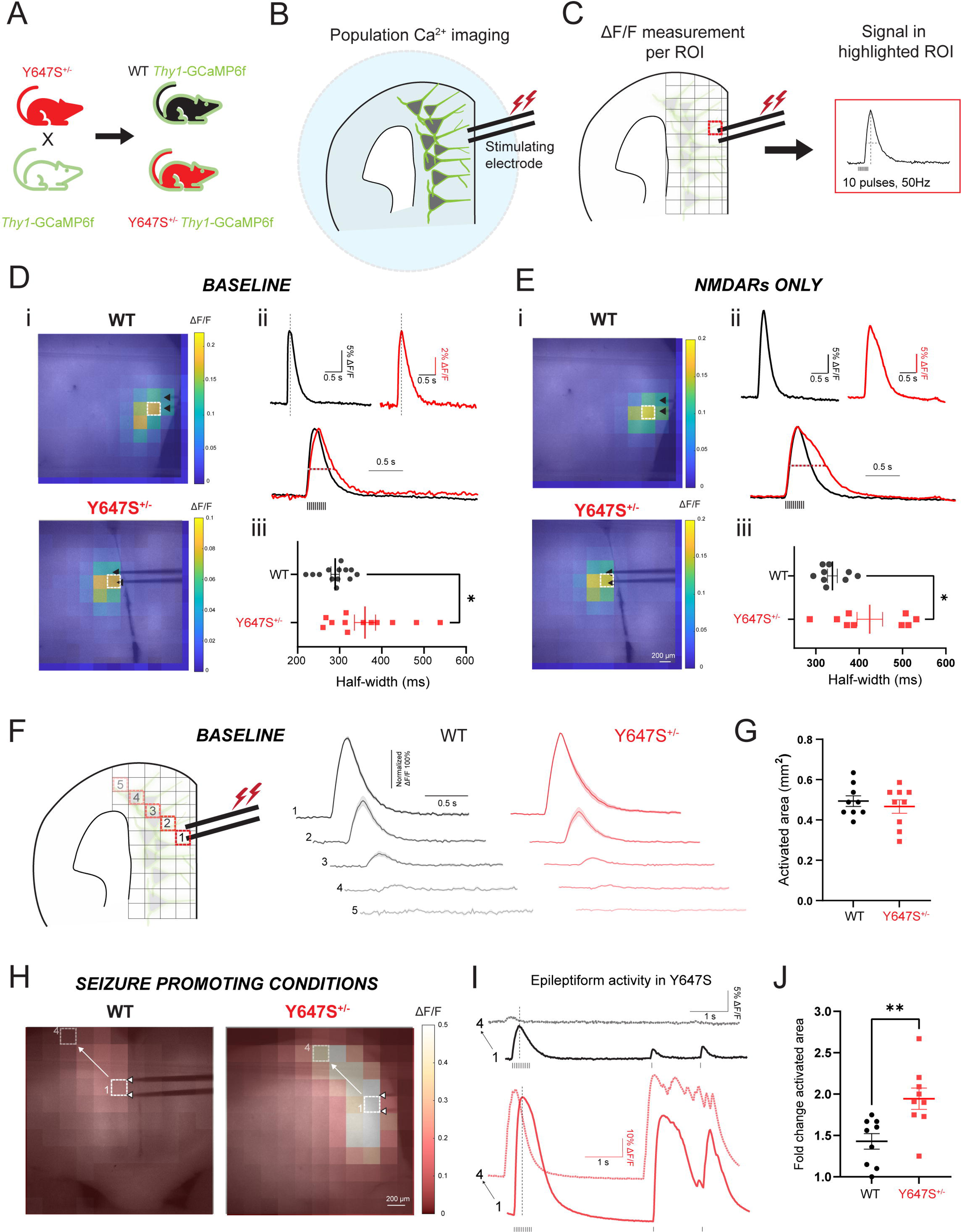
Prolonged neural population activity causes epileptiform events in *Grin1* Y647S^+/-^ brain slices. **A,** Schematic of breeding plan. Y647S^+/-^ mice were crossed with Thy1-GCaMP6f mice to generate WT and Y647S^+/-^ mice expressing GCaMP6f in layer 5 pyramidal neurons. **B-C,** Widefield calcium imaging of population neural activity in prefrontal brain slices. Stimulating electrode in the apical dendritic field of L5 neurons delivers 50Hz stimulation and the resulting fluorescence signal is captured on the widefield microscope. Change in fluorescence (ΔF/F) is calculated per 100-pixel square region (∼220 x 220 µm^2^). **D-E, i,** Heatmaps of ΔF/F at the peak signal superimposed on images of WT and Y647S^+/-^ brain slices under **D,** baseline conditions (no pharmacological blockers) and **E,** with only NMDARs active (AMPAR and GABARs blocked). Black arrows point out stimulating electrode position, white box indicates region with maximum signal. **ii,** Fluorescence signal from region highlighted in i (top). Dotted line indicates time of peak signal shown in i. Superimposed normalized responses in WT and Y647S^+/-^ are also shown (bottom). **iii,** Comparison of signal half-width in WT and Y647S^+/-^ (\**P* < 0.05, unpaired t-test). Neural population activity is prolonged in Y647S^+/-^ brain slices without any blockers at baseline (D) and with only NMDARs active (E). **F,** Spatial decay of population neural activity at baseline (no blockers) measured diagonally from the site of the stimulating electrode (left). Average fluorescence signal (normalized to the maximum) demonstrates similar spatial decay in WT and Y647S^+/-^. **G**, Total activated area of slice in WT and Y647S^+/-^. **H,** Heatmap of fluorescence signal under seizure promoting conditions in WT and Y647S^+/-^ (all GABARs blocked). Y647S^+/-^ slice shows larger activated area and greater signal. **I,** WT slice (top) shows normal decay along the diagonal and very small responses to test stimuli delivered ∼4s after the initial 50 Hz stimulation. (Bottom) Y647S^+/-^ slice exhibits reduced spatial decay and epileptiform activity with recurrent excitation triggered by the test stimuli. **J,** Fold change in total activated area of the slice from baseline to seizure-promoting conditions in WT and Y647S^+/-^ (\*\**P* < 0.01, unpaired t-test).

#### Difference in isolated and integrative NMDAR signaling but similar intrinsic properties in *Grin1* Y647S^+/-^ mice

We observed prolonged NMDAR-dependent integration and enhanced cellular NMDA currents in *Grin1* Y647S^+/-^ mice despite clearly compromised current through isolated NMDARs. This occurred in the absence of major changes in intrinsic neuronal properties except for a small but significant decrease in capacitance and increase in excitability (**Supplemental Results, Table S1, Fig S1**). This prompted us to ask how opposing loss and gain-of-function at isolated versus integrated NMDAR signaling affect cortical circuits in *Grin1* Y647S^+/-^ mice. We pursued this question using widefield calcium imaging of cortical neuronal ensembles.

### Impact on cortical circuits

#### Extended population activity triggers epileptiform events in *Grin1* Y647S^+/-^ brain slices

To assess the circuit-level consequences of the Y647S NMDAR patient variant, we performed widefield calcium imaging in brain slices from Y647S^+/-^ and littermate wildtype Thy1-GCaMP6f mice (**Fig 3A-C**, N = 4 mice per genotype). The fluorescent calcium sensor GCaMP6f is well expressed in a large population of layer 5 excitatory neurons^44^. We measured the temporal and spatial dynamics of cortical activity in response to electrically evoked glutamate release under specific pharmacological conditions.

The duration of population neural activity was significantly prolonged in Y647S^+/-^ mice compared to WT (**Fig 3D**, half-width, WT: 289 ± 10 ms, Y647S^+/-^: 360 ± 25 ms, Welch’s t test: t _(14.49)_ = 2.63, *P* = 0.019). Extended population activity in Y647S^+/-^ mice was observed in the absence of any pharmacological blockers with intact glutamatergic and GABAergic neurotransmission. Next, we pharmacologically isolated population NMDAR-mediated calcium signal using AMPAR and GABAR blockers (**Fig 3E**). Population NMDAR calcium signals were also significantly prolonged in Y647S^+/-^ brain slices (WT: 338 ± 11 ms, Y647S^+/-^: 425 ± 30 ms, t _(10.29)_ = 2.73, *P* = 0.02). We conclude that extended population neural activity is a direct circuit-level correlate of previously observed prolonged dendritic plateau potentials.

To determine whether spatial extent of cortical activity is altered in Y647S^+/-^ mice, we first measured the total activated area of the brain slice under baseline conditions with intact glutamatergic and GABAergic neurotransmission (**Fig 3F**). The total activated area of the slice was similar between WT and Y647S^+/-^ mice at baseline (**Fig 3G**, WT: 0.49 ± 0.03 mm^2^, Y647S^+/-^ : 0.46 ± 0.03 mm^2^, t_(16)_ = 0.63, *P* = 0.53). However, under seizure-promoting conditions where GABAergic transmission was blocked, Y647S^+/-^ brain slices exhibited widespread epileptiform activity characterized by recurrent and persistent excitation even in the absence of stimulation (**Fig 3H-I**). This resulted in larger area of the slice being activated in Y647S^+/-^ (**Fig 3J**, 1.9 ± 0.1-fold change from baseline) compared to WT (1.4 ± 0.1, t _(16)_ = 3.24, *P* = 0.005).

We conclude that the Y647S mutation results in prolonged cortical network excitation that can transition into widespread and persistent epileptiform activity. Our circuit-level characterization of a *GRIN1* patient variant provides the first *ex vivo* proof of epileptiform activity and the conditions that can initiate seizures in mice and may do so in patients.

### Negative feedback links loss of isolated NMDAR function to extended dendritic excitation

Our multi-scale investigation revealed deficient current through isolated synaptic NMDARs, but paradoxically extended NMDAR integration that prolongs network excitation resulting in epileptiform activity (**Fig 4**). We sought to identify the mechanism which results in seizure-promoting excitation despite a major reduction in isolated NMDAR function. Such a mechanism would enable precision intervention to prevent seizures in Y647S^+/-^ mice. One candidate mechanism is Ca^2+^-dependent negative feedback of NMDARs: Ca^2+^ influx through the NMDAR activates small conductance SK calcium-activated potassium channels which hyperpolarize the membrane, restoring NMDAR Mg^2+^ block and terminating NMDA receptor activation^33,45^. In Y647S^+/-^ neurons, reduced NMDAR Ca^2+^ influx could result in insufficient activation of SK channels. The consequence of this “brake failure” would be prolonged dendritic plateau potentials^46^ and enhanced whole-cell NMDA currents. We tested this hypothesis by manipulating the two arms of this negative feedback mechanism: i) SK channels and ii) Mg^2+^ block of NMDARs in Y647S^+/-^ mice.

**Figure 4.**
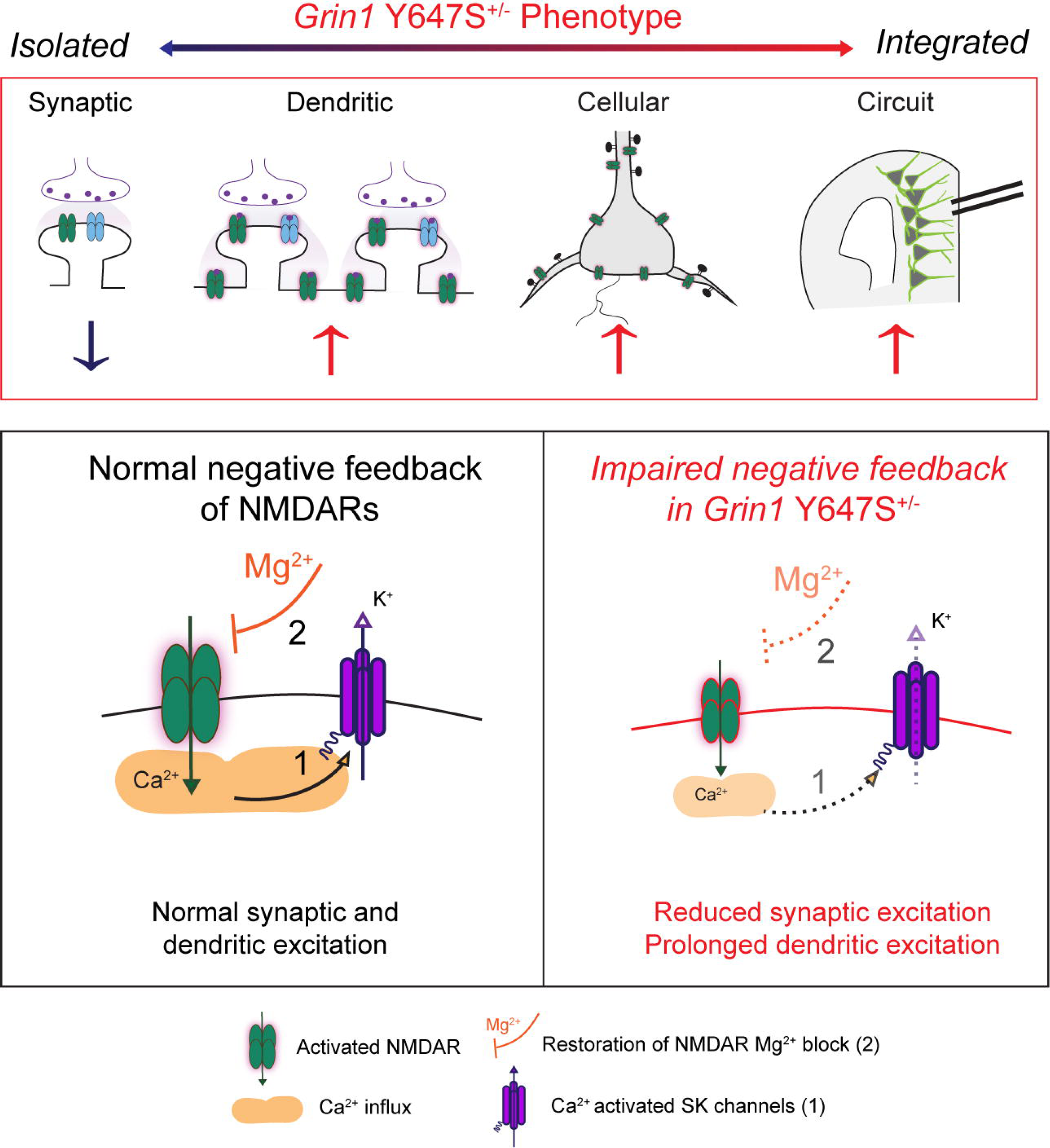
Negative feedback mechanism links loss of isolated NMDAR function to extended dendritic excitation. (Top) Summary schematic of results shows reduced isolated receptor-level currents, but prolonged NMDAR-dependent integration, increased whole-cell NMDA currents, and prolonged circuit excitation in *Grin1* Y647S^+/-^ mice. (Bottom) Potential mechanism. (Left) Normal Ca^2+^ influx through WT NMDARs activates small conductance SK_Ca_ potassium channels (1) which hyperpolarize the membrane, restoring NMDAR Mg^2+^ block (2), terminating NMDAR activation. (Right) Reduced NMDAR Ca^2+^ influx through Y647S^+/-^ NMDARs causes insufficient activation of SK channels, with impaired negative feedback of NMDARs. This “brake failure” can prolong dendritic and circuit excitation, promoting seizures.

### Boosting negative feedback: Potentiating SK channels prevents extended NMDAR integration in *Grin1* Y647S^+/-^

NMDARs typically receive powerful local feedback from Ca^2+^ activated potassium channels, including SK family members, whose hyperpolarizing currents inactivate NMDARs by restoring Mg^2+^ block^33,46^. Insufficient activation of SK channels could cause prolonged NMDAR-dependent integration in Y647S^+/-^ neurons. To test this hypothesis, we assessed whether the highly potent SK channel activator NS309 could reduce plateau potential duration in Y647S^+/-^ neurons to WT levels. NS309 increases the calcium sensitivity of SK channels^47,48^, enabling their activation even with insufficient Ca^2+^ influx in Y647S^+/-^ neurons (**Fig 5A**). NS309 successfully eliminated the prolonged depolarization in Y647S^+/-^ neurons (**Fig 5B**, N = 4, 6 mice for WT and Y647S), restoring the duration of plateau potentials to WT levels (**Fig 5C**, Plateau duration at highest stimulus: WT = 0.39 ± 0.04 s, Y647S^+/-^ = 0.88 ± 0.09 s, Y647S^+/-^ + NS309 = 0.48 ± 0.14 s).

**Figure 5.**
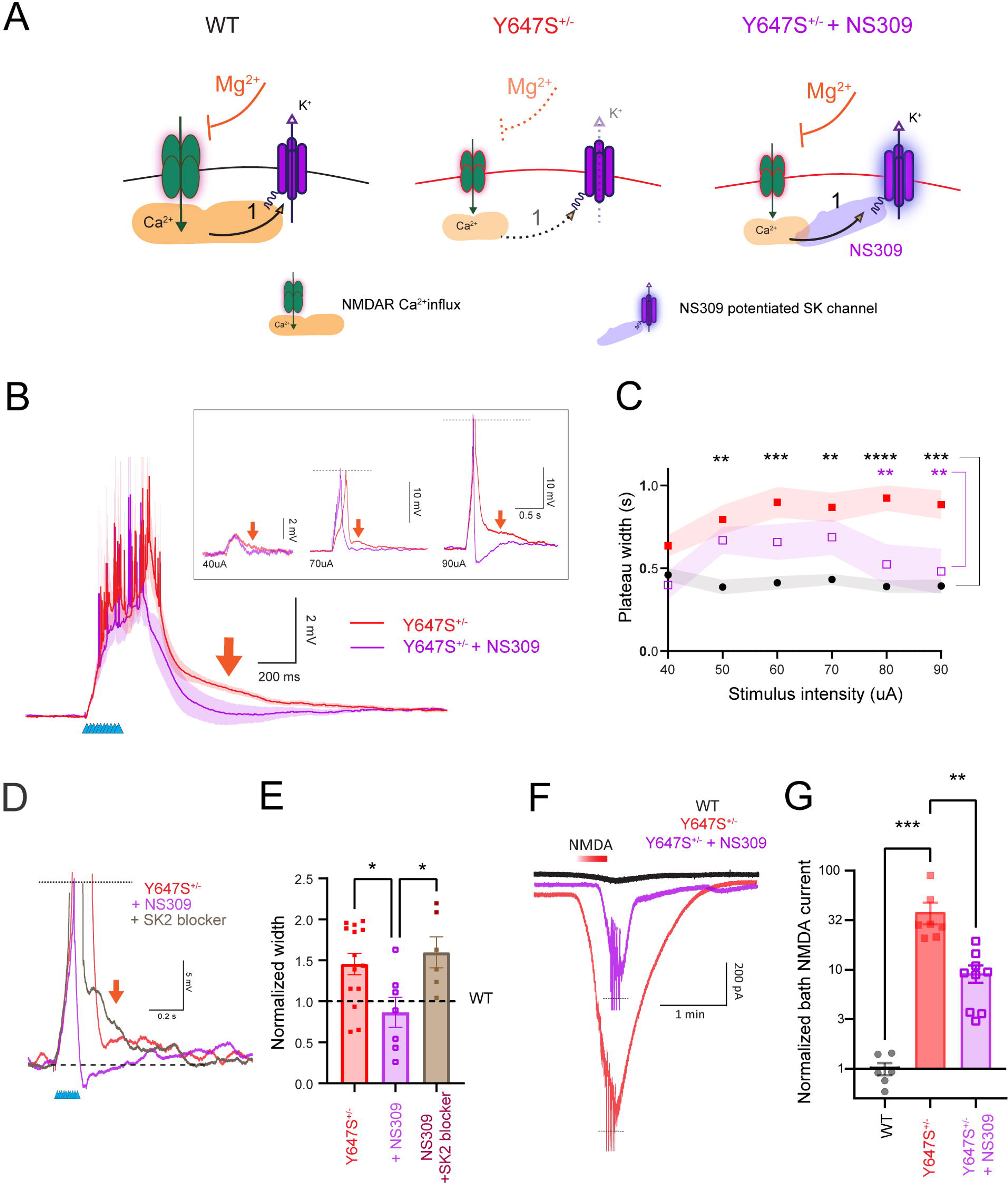
Potentiating SK channels prevents extended NMDAR integration in *Grin1* Y647S^+/-^ mice. **A,** Schematic of potential mechanism showing impaired negative feedback in Y647S^+/-^ neurons due to insufficient Ca^2+^ influx compared to WT and the effect of NS309 in boosting SK channel Ca^2+^ sensitivity, thereby restoring negative feedback in Y647S^+/-^ neurons. **B,** Average NMDAR plateau potential in Y647S^+/-^ neurons at 70 µA with extended tail indicated by red arrow. Application of 10 µM NS309 to the slice restores normal duration and terminates the NMDAR plateau potential in Y647S^+/-^ (Y647S^+/-^ + NS309). (Inset) Restoration of plateau potential duration by NS309 at increasing stimulus intensities in a Y647S^+/-^ neuron. **C,** Total width of the NMDAR plateau potential in WT, Y647S^+/-^, and Y647S^+/-^+ NS309 neurons (\*\**P* < 0.01, \*\*\**P* < 0.001, \*\*\*\**P* < 10^-4^, Tukey’s post hoc. Black stars: WT vs Y647S^+/-^, purple stars: Y647S^+/-^ vs Y647S^+/-^ + NS309). **D**, SK2 channel blocker (Leidab7, 100 nM) prevents NS309 from reducing plateau potential duration in Y647S^+/-^ neurons. **E,** Normalized NMDAR plateau width in Y647S^+/-^ neurons with the addition of NS309 and NS309+SK2 blockers (\**P* < 0.05, Sidak’s post hoc). **F,** Whole-cell NMDA current evoked by 20 µM NMDA in WT, Y647S^+/-^, and Y647S^+/-^ + NS309 neurons. **G,** Peak whole-cell NMDA current normalized to WT in Y647S^+/-^ and Y647S^+/-^ + NS309. (\*\**P* < 0.01, \*\*\**P* < 0.001, Sidak’s post hoc).

NS309 can act on both SK2 and SK3 subtypes of SK channels in addition to intermediate conductance IK channels^47^. To determine the channel subtype involved in NS309-restoration of plateau potential duration, we tested whether SK2 channels were the target by coapplying the SK2 blocker Leidab7. SK2 blockers occluded the effect of NS309, preventing the reduction in plateau potential duration in Y647S^+/-^ neurons (**Fig 5D**).Normalized plateau potential duration in Y647S^+/-^ (1.46 ± 0.13) was significantly reduced by NS309 (0.87 ± 0.18, t _(32)_ = 2.88, *P* = 0.028), but this decrease was prevented by the co-application of SK2 blockers (1.59 ± 0.19, t _(32)_ = 2.97, *P* = 0.022, **Fig 5E**). We conclude that the potentiation of SK2 channels by NS309 prevents prolonged dendritic excitation in Y647S^+/-^ neurons by restoring NMDAR negative feedback.

Next, we examined whether NS309 would reduce the impact of the Y647S mutation on whole-cell NMDA currents (**Fig 5F**). NS309 caused a 76% reduction (**Fig 5G**, t _(19)_ = 3.98, *P* = 0.002) of whole-cell NMDA currents in Y647S^+/-^ neurons which typically exhibit higher NMDA currents compared to WT (t _(19)_ = 4.63, *P* = 5 * 10^-4^). These experiments established that boosting negative feedback via SK channels was sufficient to prevent prolonged integration and enhanced cellular NMDA currents in Y647S^+/-^ neurons. We infer that impaired NMDAR negative feedback underlies seizure promoting excitation in Y647S^+/-^ mice. We further tested this conclusion by evaluating whether deliberately reducing NMDAR negative feedback would disproportionately impact Y647S^+/-^ neurons.

### Reducing negative feedback: Lowering extracellular magnesium promotes NMDAR hyperexcitation in *Grin1* Y647S^+/-^

We directly evaluated NMDAR-negative feedback in Y647S^+/-^ neurons by altering extracellular magnesium levels. If inadequate SK channel recruitment reduces NMDAR Mg^2+^ block in Y647S^+/-^ neurons, we predicted that they would be highly sensitive to reductions in extracellular magnesium levels. Even a small reduction in extracellular Mg^2+^ would cause disproportionate NMDAR excitation in Y647S^+/-^ neurons while WT neurons would not be affected (**Fig 6A**).

**Figure 6.**
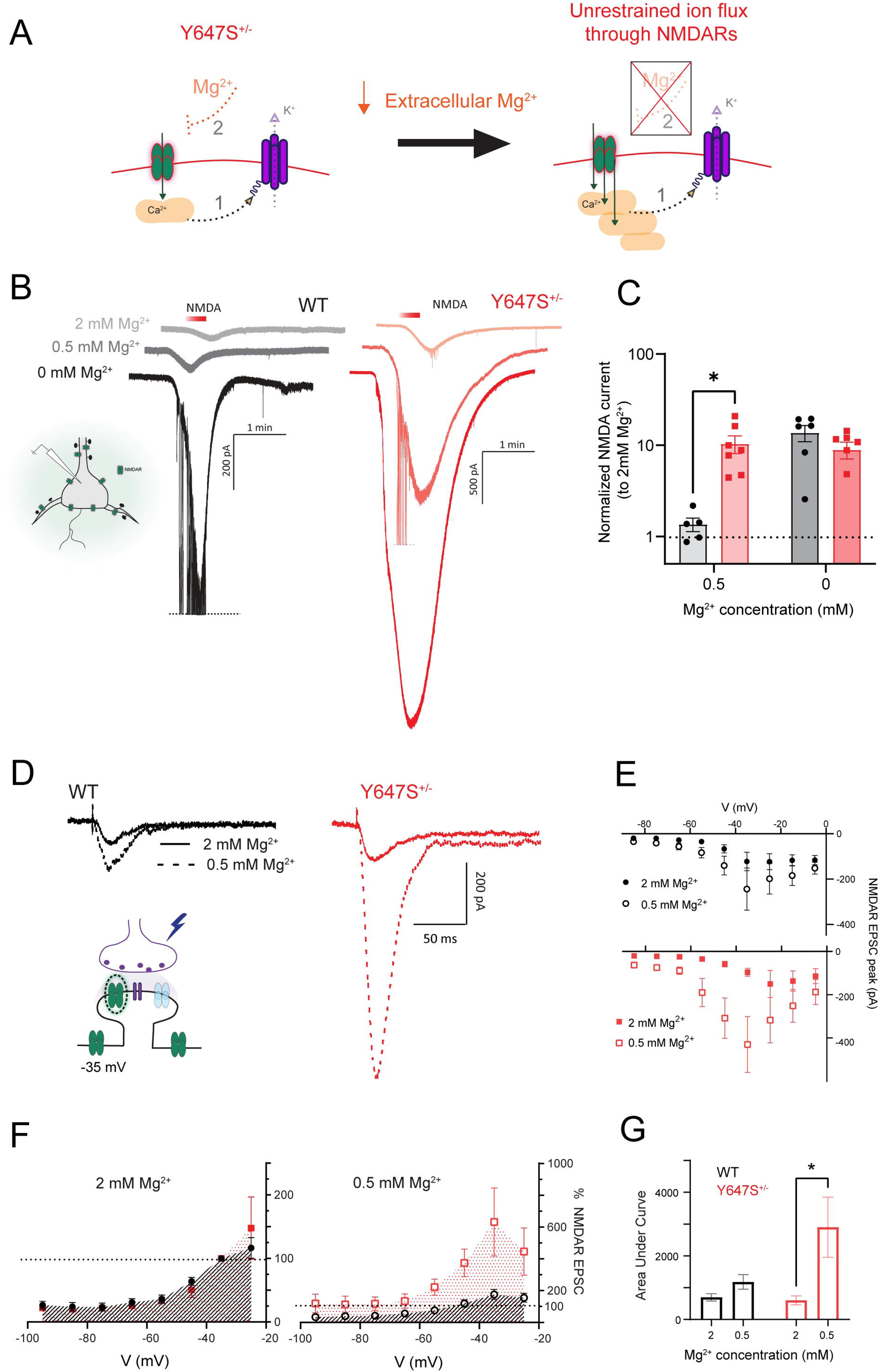
Reducing extracellular magnesium causes disproportionate NMDAR hyperexcitation in *Grin1* Y647S^+/-^ mice. **A,** Schematic illustrating the consequence of reducing extracellular Mg^2+^ levels in Y647S^+/-^ neurons. Due to impaired SK-channel mediated negative feedback, this can cause unrestrained NMDAR activation. **B,** Whole-cell NMDA current evoked by 20 µM NMDA in a WT and Y647S^+/-^ neuron at 2, 0.5, and 0 mM extracellular Mg^2+^ concentrations. **C,** Peak NMDA current at 0.5 and 0 mM Mg^2+^ in WT and Y647S^+/-^ neurons normalized to the peak current at 2 mM Mg^2+^ within each genotype (\**P* < 0.05, Sidak’s post hoc). Y647S^+/-^ NMDA current is disproportionately increased by reducing Mg^2+^ from 2 to 0.5 mM. **D,** Electrically-evoked NMDAR EPSCs measured at holding potential of -35 mV in WT and Y647S^+/-^ neurons at 2 and 0.5 mM extracellular Mg^2+^ concentration. Potassium gluconate patch pipettes containing 5 mM QX314 were used to assess integrated NMDAR signaling with active SK channels. **E**, Average current-voltage relationship (I-V curve) between NMDAR EPSC amplitude and membrane potential at 2 and 0.5 mM Mg^2+^ in WT (top) and Y647S^+/-^ (bottom). **F,** Normalized NMDAR activation curve (I-V curve normalized to peak at -35 mV, 2mM Mg^2+^) in each genotype at 2 mM (left) and 0.5 mM (right) extracellular Mg^2+^ concentrations. **G,** Area under the normalized NMDAR activation curve at 2 and 0.5 mM Mg^2+^ in WT and Y647S^+/-^ neurons (\**P* < 0.05, Sidak’s post hoc).

We tested this first by measuring whole-cell NMDA currents at different Mg^2+^ concentrations. Y647S^+/-^ neurons showed a 10-fold increase in the amplitude of NMDA currents measured at 0.5 mM Mg^2+^ compared to 2 mM Mg^2+^ (Welch’s t test: t _(6.15)_ = 3.58, *P* = 0.01). In contrast, NMDA currents in WT neurons did not differ majorly between 2 mM Mg^2+^ and 0.5 mM Mg^2+^ (t _(12)_ = 1.67, *P* = 0.12). The fold-change in NMDA current from 2 to 0.5 mM Mg^2+^ was much larger in Y647S^+/-^ (10.4 ± 2.3, t _(21)_ = 2.89, *P* = 0.018) compared to WT (1.4 ± 0.2) (**Fig 6B – C**, 2-way ANOVA, genotype x Mg^2+^: F _(1, 21)_ = 10.24, *P* = 0.004). Complete removal of extracellular Mg^2+^ from 2 to 0 mM caused a similar fold-change in NMDA currents in both WT (13.7 ± 2.8) and Y647S^+/-^ (8.9 ± 1.9, Sidak’s post hoc: t _(21)_ = 1.6, *P* = 0.23). Y647S^+/-^ neurons show disproportionate NMDAR excitation with small reductions in extracellular Mg^2+^ due to inadequate negative feedback, whereas complete removal of Mg^2+^ equivalently eliminates negative feedback in both genotypes.

Next, we tested whether NMDARs activated by endogenous glutamate release in Y647S^+/-^ neurons were also disproportionately impacted by reduced extracellular Mg^2+^. To retain SK channel contribution, we used a modified patch solution with QX-314 (**Supplemental Methods**) and measured NMDAR EPSCs at different membrane potentials. The current-voltage relationship (I-V) of WT and Y647S^+/-^ NMDARs was identical at 2 mM extracellular Mg^2+^. NMDAR EPSCs showed the expected increase at positive membrane potentials as Mg^2+^ block is relieved, reaching peak amplitude at -35 mV in both genotypes (**Fig 6F**). Lowering Mg^2+^ from 2 to 0.5 mM caused a disproportionate increase in EPSC amplitude in Y647S^+/-^ neurons (6.3 ± 2.1-fold-change at -35 mV) compared to WT (1.8 ± 0.3, t _(13)_ = 2.26, *P* = 0.04, **Fig 6D-E**). Normalized NMDAR activation curves are identical in WT and Y647S^+/-^ at 2 mM Mg^2+^ (Area under curve: t _(26)_=0.14, *P*= 0.98) but are significantly enhanced in Y647S^+/-^ neurons at 0.5 mM Mg^2+^ (t _(26)_=2.61, *P*= 0.03, **Fig 6F-G**). The total area under the curve is significantly increased at 0.5 compared to 2mM Mg^2+^ in Y647S^+/-^ (t _(13)_ = 3.33, *P* = 0.011), but not in WT neurons (t _(13)_ = 0.75, *P* = 0.71; **Fig 6G**; 2-way RM ANOVA, Mg^2+^ concentration: F _(1, 13)_ = 8.691, *P* = 0.011). Both whole-cell NMDA currents and endogenous NMDAR responses show that Y647S^+/-^ neurons are specifically vulnerable to NMDAR hyperexcitation when extracellular Mg^2+^ is reduced.

Bidirectional manipulation of NMDAR negative feedback, first by boosting SK channel activation, next by reducing extracellular Mg^2+^ levels revealed impaired negative feedback in Y647S^+/-^. This results in protracted integrated NMDAR signaling that promote seizures in Y647S^+/-^ mice despite the initial loss of isolated NMDAR currents. Therefore, we conclude that boosting NMDAR negative feedback is a promising approach to treat NMDAR hyperactivation and prevent seizures in Y647S^+/-^ mice.

### Magnesium threonate treatment in vivo successfully treats seizures in *Grin1* Y647S^+/-^ mice

To boost NMDAR negative feedback in vivo and prevent seizures in Y647S^+/-^ mice, we aimed to increase brain levels of Mg^2+^. Magnesium supplementation has been historically used to treat seizures occurring during pre-eclampsia^49,50^ and other forms of epilepsy^51^. However, previous formulations like magnesium sulphate do not achieve effective Mg^2+^ increase in the brain and require invasive administration^52^. We opted for Magnesium L-Threonate, a recent formulation that can be administered orally to increase brain levels of Mg^2+^ with pro-cognitive effects^34,53,54^. Adult littermate WT and Y647S^+/-^ mice of both sexes were given normal drinking water (control) or 0.5 % w/v magnesium threonate (MgT) with *ad libitum* access for 12 weeks. We allowed for 6 weeks of pre-treatment before assessing seizures to ensure efficacy well beyond the “honeymoon period” that is typical for antiepileptic drugs^55,56^.

At 7 weeks, 83% of untreated control Y647S^+/-^ mice (5/6) displayed prominent behavioural seizures upon handling, characterized by loss of posture, body and facial convulsions. Remarkably, none of the MgT treated Y647S^+/-^ mice (0/7) had seizures (Fisher’s exact test: *P* = 0.005, **Fig 7A**). Seizures in control Y647S^+/-^ mice lasted up to 1.7 minutes and were quite severe (mean: 41 ± 15 s, compared MgT treated Y647S^+/-^: t _(11)_ = 3.04, *P* = 0.011). These involved loss of posture, facial convulsions, limb clonus, and body twitching that were quantified by our modified Racine scale (RS), with 3/6 mice showing highest severity (RS 5). In contrast, MgT treated Y647S^+/-^ mice did not display seizures of any severity (unpaired t-test: t _(11)_ = 3.95, *P* = 0.002; **Fig 7B**). MgT appears to show 100% efficacy in preventing the spontaneous handling-induced behavioural seizures at 6-7 weeks of treatment. In addition, we observed MgT to be beneficial in improving weight gain and reducing hyperactivity in Y647S^+/-^ mice (**Supplemental Fig S2**).

**Figure 7.**
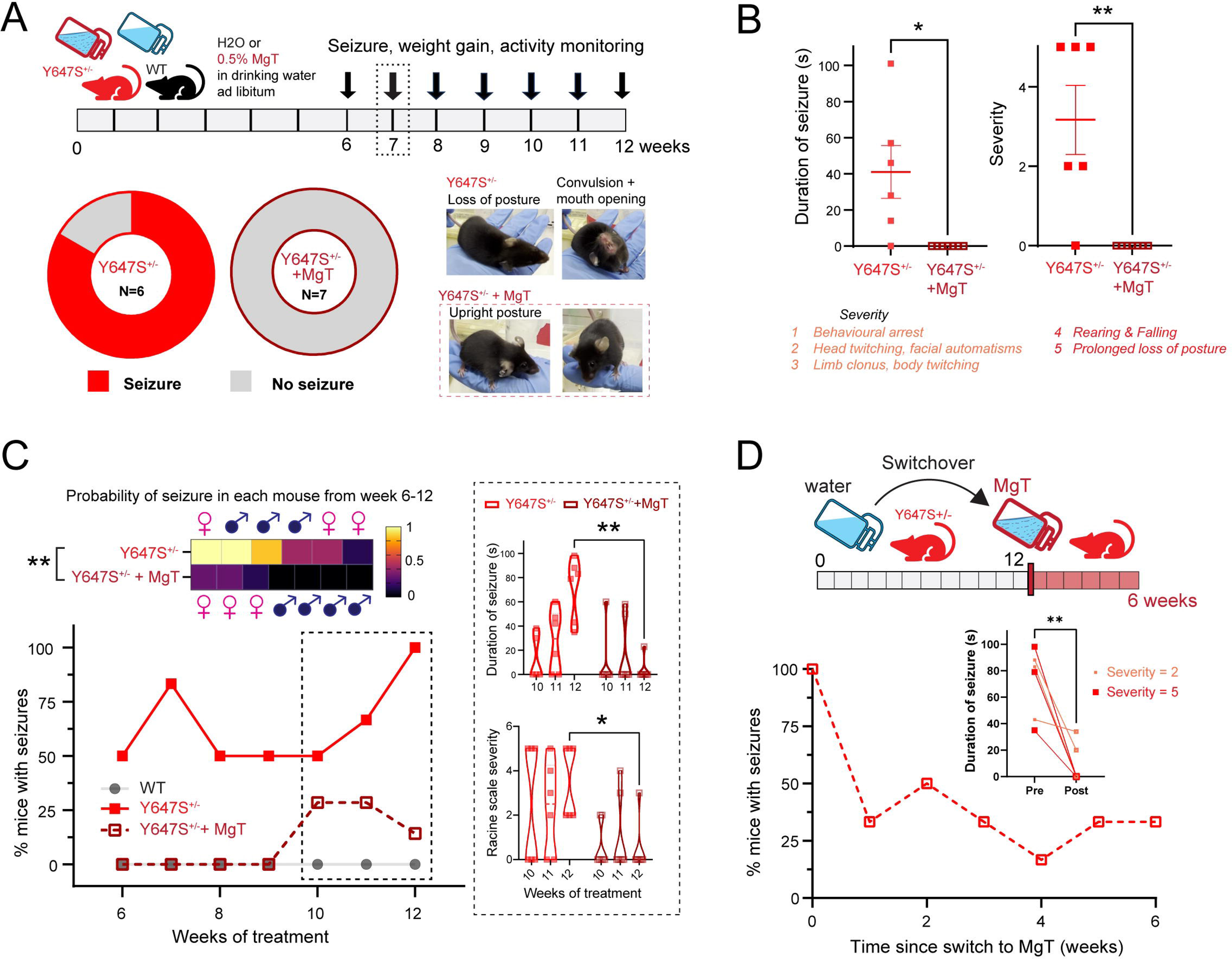
Magnesium threonate treatment *in vivo* successfully treats seizures in *Grin1* Y647S^+/-^ mice. **A,** Schematic of experimental plan. Adult WT and littermate Y647S^+/-^ mice were given *ad libitum* access to drinking water with 0.5% magnesium-L-threonate (MgT) or normal water (controls) for 12 weeks, with weekly monitoring of handling induced seizures and weight gain from 6 weeks onwards. Donut charts show proportion of control Y647S^+/-^ (5/6) and Y647S^+/-^ + MgT (0/7) mice exhibiting handling induced seizures at week 7 of treatment. Inset pictures show example convulsions with loss of posture and mouth opening in untreated Y647S^+/-^ mouse (top) versus normal upright posture in Y647S^+/-^ + MgT mouse. **B,** Duration of seizures (left) and Severity on our modified Racine scale (right) in Y647S^+/-^ and Y647S^+/-^ + MgT mice at week 7 (\**P* < 0.05, \*\**P* < 0.01, unpaired t-test). **C,** Seizure incidence and severity for the whole duration of the treatment. Top: Average seizure probability per mouse from 6-12 weeks shown as a heatmap, with sexes of individual mice shown above and below (\*\**P* <0.01, unpaired t-test). Bottom: Percentage of mice with handling induced seizures from weeks 6-12 of the treatment period. (Inset) Duration (top) and severity of seizures (bottom) in control and MgT treated Y647S^+/-^ mice for weeks 10-12 (\**P* <0.05, \*\**P* < 0.01, Sidak’s post hoc). MgT remains effective in suppressing seizures until 12 weeks. **D,** Top: Experimental schematic, untreated Y647S^+/-^ from the same cohort as C were switched to MgT treatment at 12 weeks. Bottom: percentage of mice with seizures for 6 weeks from the switch. (Inset) Duration of seizures at week 0 (before switching to MgT) and at 6 weeks after switching to MgT (light red indicates mild seizures with severity 2, dark red corresponds to severe convulsive seizures, \*\**P* < 0.01, paired t-test). Severe seizures are eliminated after switching to MgT.

We examined the long-term efficacy of MgT by monitoring the percentage of mice exhibiting seizures up to 12 weeks of treatment (**Fig 7C**). Seizures in control Y647S^+/-^ mice became more frequent, with all mice developing seizures by the 12^th^ week, and 2/6 mice having seizures every single week. In contrast, MgT-treated mice showed no signs of seizures until 10 weeks, but a proportion (3/7) started to exhibit breakthrough seizures in weeks 10-12. The duration of seizures in weeks 10-12 was significantly reduced in MgT-treated Y647S^+/-^ mice compared to untreated control Y647S^+/-^ mice (2-way ANOVA, week x treatment: F _(2, 22)_ = 8.28, *P* = 0.002; Sidak’s post hoc: t _(5.98)_ = 6.16, *P* = 0.003). MgT-treated Y647S^+/-^ mice also had less severe seizures compared to untreated controls (2-way ANOVA, Treatment: F _(1, 11)_ = 5.763, *P* = 0.03; Sidak’s post hoc at week 12: t _(33)_ = 3.11, *P* = 0.011). Breakthrough seizures in MgT-treated mice all occurred in female mice (n = 3/7, **Fig 7C**) indicating that females might be more vulnerable to reductions in treatment efficacy. Overall, the average seizure probability was significantly higher in control Y647S^+/-^ mice (0.59 ± 0.14) compared to MgT treated Y647S^+/-^ mice (0.11 ± 0.05, unpaired t-test: t _(11)_ = 3.36, *P* = 0.034).

Finally, we examined whether acute treatment with MgT could be effective in the untreated Y647S^+/-^ mice that had developed severe seizures (**Fig 7D**). This experiment was critical to determine whether treatment in adulthood after full development of disease pathology could still be effective. Control Y647S^+/-^ mice that were switched to MgT treatment at week 12 showed an immediate reduction in seizure occurrence with complete elimination of severe seizures after two weeks of treatment. The duration of seizures was significantly reduced in all mice after 6 weeks of MgT treatment (Paired t-test: t _(5)_ = 4.54, *P* = 0.006). We conclude that MgT intervention is acutely effective in reducing seizure occurrence and severity in *Grin1* Y647S^+/-^ mice.

Our work reveals a novel mechanism behind epilepsy caused by GluN1 patient variant NMDA receptors. We show that loss-of-function of NMDA receptors can result in prolonged cortical excitation that promotes seizures due to deficient negative feedback within neurons. Boosting NMDAR negative feedback with an oral magnesium supplement successfully treats seizures. These results confirm that focusing on isolated NMDAR consequences can be misleading and evaluating NMDARs in multiple functional contexts is essential to predict effective treatments for *GRIN* disorders.

## Discussion

Despite the diverse functional contexts of NMDARs, NMDAR patient variants have mainly been studied at the level of isolated receptors. Our work in *Grin1* Y647S^+/-^ mice reveals opposing consequences on isolated versus integrative NMDAR signaling, leading to an unexpected seizure treatment.

### Reconciling loss/gain of function of NMDAR patient variants with multi-scale assessment

The GluN1 Y647S variant is predicted to be loss-of-function ^8,20,57^ based on decreased currents and surface expression in oocytes^20^, HEK cells, and neurons^31^. But there are also reports of increased agonist affinity in Y647S and Y647C GluN1 variant NMDARs^8,58^. Previous work has shown that severe knockdown of the GluN1 subunit has subcellular compartment-specific consequences^28^, prompting a multi-scale assessment in *Grin1* Y647S^+/-^ mice. We found reduced current through isolated NMDARs, consistent with receptor loss-of-function. However, integrated and circuit-level NMDAR signaling were greatly enhanced, promoting seizure vulnerability. These opposing effects show that functional context dictates NMDAR deficits and receptor-level impact is insufficient to characterize patient variants.

There is growing recognition that integrative parameters are necessary to understand glutamate receptor patient variants^14,59–61^. Yet there is a bias towards receptor-level impact when classifying ion channel variants, especially in emerging machine learning techniques ^62,63^. While receptor-level impact is a first step to predict variant pathogenicity, functional context is essential to determine overall phenotype and predict treatments.

### Dendritic plateau potentials are key to integrated assessment of NMDAR variants

Plateau potentials occur in thin dendrites of pyramidal neurons when extrasynaptic NMDARs are activated by glutamate spillover^43,64,65^. Plateau potentials are linked to cognitive integration^66^, synaptic plasticity^67^, and enable the formation of neuronal ensembles^65,68^. We found severely prolonged plateau potentials in *Grin1* Y647S^+/-^ neurons, with plateau duration serving as a biomarker of seizure-promoting network excitation. Plateau potential properties are linked to dendritic calcium-activated potassium channels (SK, BK)^46,69^ and are known to alter excitability in Fragile X^70,71^ and Dravet syndrome^72^.

NMDAR Ca^2+^ influx activates potassium channels that provide essential negative feedback by helping to restore NMDAR magnesium block^73,74^. High-affinity SK channels on dendritic shafts and synapses exert powerful inhibition of NMDAR postsynaptic potentials^33,46,75,76^. We find that enhancing calcium sensitivity of SK channels in Y647S^+/-^ mice restored normal plateau potential duration. In contrast, reducing negative feedback by lowering extracellular Mg^2+^ levels caused disproportionate NMDAR hyperexcitation in Y647S^+/-^. While one might suspect directly reduced efficacy of Mg^2+^ block in Y647S^+/-^ NMDARs, the similarity in current-voltage (I-V) curves of WT and Y647S^+/-^ NMDARs at physiological Mg^2+^ concentration indicate that is not the case. Moreover, HEK cell results indicated intact Mg^2+^ block of isolated NMDARs^29,58^. We conclude that insufficient negative feedback of impaired NMDARs constitutes a cell-autonomous mechanism that increases seizure vulnerability.

### Targeting NMDAR negative feedback to treat seizures caused by GRIN variants

Treatments for *GRIN* disorder have previously focused on receptor-level impact. NMDAR antagonists ketamine and memantine reduced seizure burden in a GluN1 gain-of-function patient^9^, and glutamate receptor antagonists radiprodil^77,78^, perampanel^79,80^ are effective for other gain-of-function variants. L-serine, a precursor of NMDAR co-agonist D-serine, improves motor and cognitive functions with limited effect on seizures in loss-of-function variants^81–83^. However, seizures in loss-of-function GluN1 variants may arise from compensatory processes downstream of the receptor due to its obligatory nature. We show that indirect loss of NMDAR negative feedback underlies seizure vulnerability in the GluN1 Y647S variant. Instead of focusing on the isolated receptor, we chose to boost negative feedback using magnesium supplementation to target integrated NMDAR signaling.

Magnesium sulphate treatment is well-supported for seizures in preeclampsia^49^ and other forms of epilepsy^51,84^. However, it has been difficult with previous Mg^2+^ formulations to increase brain levels of Mg^2+^ efficiently. Intrathecal magnesium sulphate reduced seizures in a *GRIN1* patient with receptor gain-of-function^52^, but this was a highly invasive procedure. Magnesium threonate (MgT) is better transported into the brain with oral administration^34^ and benefits brain health and cognition^85,86^. Here, oral MgT treatment remarkably reduced the incidence and severity of seizures in *Grin1* Y647S^+/-^ mice chronically over 3 months, and acutely within 2 weeks. The average western diet is deficient in magnesium^87^ and severe hypomagnesemia causes seizures^88^. Genetic variants in magnesium transporters are also linked to risk of several neuropsychiatric illnesses^89^. It is critical to evaluate and treat *GRIN* disorders against the backdrop of common dietary and genetic vulnerability in brain magnesium levels.

### Caveats and future directions

We highlight the general principle that changes in isolated NMDARs can be misleading, with opposing loss and gain-of-function at different contexts of neural signaling. While we limited our examination to principal neurons of the cortex, the integrative NMDAR phenotype we identified was consistent at the circuit-level and predicted an effective *in vivo* treatment. In brain slices ex vivo, boosting Ca^2+^ sensitivity of SK channels acts in a similar manner to increasing Mg^2+^ levels; both help put the "brakes" on integrative NMDAR signaling. While we focused on reducing seizures in *Grin1* Y647S^+/-^ mice with MgT treatment, other changes in synaptic plasticity^29^ and neuronal morphology may contribute to cognitive deficits. These effects are not easily separable from the detrimental cognitive burden and long-term effects of severe seizures. A precision treatment for seizures is an essential first step to benefit future assessment of cognitive function and other neurological changes in *GRIN* disorder.

### Significance

Using mice with a patient-variant in the obligate GluN1 NMDAR subunit, we show that seizure-promoting excitation arises from loss-of-function via impaired negative feedback. We successfully target this unexpected mechanism to treat seizures, demonstrating the importance of functional context to decipher and treat NMDAR dysfunction in *GRIN* disorder.

## Data availability

All experimental data generated in this paper is available upon reasonable request to the corresponding author.

## Acknowledgements

We thank Wendy Horsfall for expert technical assistance.

## Supplemental Material

Supplemental material includes Supplemental Methods and Results, Supplemental Table S1, Supplemental Figures S1-S2 and is attached as a separate PDF.

