## Supplemental Material for "Context-dependent NMDA receptor dysfunction predicts seizure treatment in mice with human GluN1 variant"

### Supplementary Material

- Suppl. Methods
- Suppl. Results
- Suppl. Table S1
- Suppl. Figure S1
- Suppl. Figure S2
- References

### Supplemental Methods

Adult male and female *Grin1* Y647S$^{+/-}$ (153 ± 12 days old) and littermate wildtype (WT) mice (160 ± 13 days old) were used for slice electrophysiology experiments.

#### Patch solutions

*Regular patch solution (K-gluconate):* in mM, 120 potassium gluconate, 5 KCl, 10 HEPES, 2 $MgCl_2$, 4 $K_2$-ATP, 0.4 $Na_2$-GTP, and 10 sodium phosphocreatine, pH 7.3. Unless otherwise specified, K-gluconate patch solution was used for most experiments, with the sodium channel blocker QX-314 (5 mM) added in a subset of specified experiments.
*Cesium gluconate patch solution (Cs-gluconate):* in mM, 105 Cesium gluconate, 17.5 CsCl, 10 HEPES, 10 BAPTA Tetracesium salt, 2 Mg-ATP, 0.3 $Na_2$-GTP, 5 QX-314 bromide, pH adjusted to 7.3 with CsOH.

#### Protocols for multi-scale NMDAR assessment

1. <u>*Evoked excitatory postsynaptic currents*</u>

   α-amino-3-hydroxy-5-methyl-4-isoxazolepropionic acid receptor (AMPAR) and NMDAR-mediated evoked excitatory postsynaptic currents (EPSCs) were measured using Cs-gluconate patch pipettes in voltage-clamp at a holding potential of −60 and +40 mV respectively. Cs-gluconate pipettes allowed us to isolate currents through the glutamate receptors by blocking all other sodium, potassium and calcium ion channels that might get activated at more positive membrane potentials. A bipolar stimulating electrode (FHC) in the basal dendritic field ~100 µm from the soma of the recorded layer 5 pyramidal neuron was used to stimulate synaptic glutamate release with single pulses of 40 µs duration delivered at 0.1 Hz. To calculate synaptic AMPA/NMDA ratios, the NMDAR component of the EPSC at +40 mV was taken to be the amplitude at 3 * $tau_{decay}$ of the AMPAR EPSC at -60mV. NMDAR EPSC half-width was measured after normalizing the amplitudes of all cells. Gamma-aminobutyric acid receptor (GABAR) blockers (Picrotoxin 20 µM, *Alomone Labs*, CGP52432 1 µM, *Tocris*) were included in all recordings, and AMPAR blocker (CNQX or NBQX 20 µM, *Alomone Labs*) was used to isolate synaptic NMDAR currents at +40 mV. Miniature EPSCs (mEPSC) were measured in the presence of sodium channel antagonist tetrodotoxin (TTX, 2µM) to examine presynaptic release probability and postsynaptic strength.

2. <u>*Dendritic NMDAR plateau potentials*</u>

   To measure the impact of the Y647S mutation on NMDAR-dependent dendritic integration, we evoked basal dendritic plateau potentials[1–3], by delivering 10 pulses of 50 Hz stimuli in the presence of AMPAR and GABAR blockers (Picrotoxin 20 µM, CGP52432 1 µM, NBQX 20 µM). Current clamp recordings at -75 mV were obtained using K-gluconate patch solution. Amplitude and duration of NMDAR plateau potentials was measured using Axograph and Clampfit 10.2 (Molecular Devices). The peak amplitude of NMDAR plateau potentials was capped at the spiking threshold, and peaks were normalized to the average spike threshold for comparison across stimulus intensities and genotypes. The total width of the plateau potential, and area under the plateau potential for 1s post-stimulation (late area) were used to assess the duration of the plateau potentials. D-APV, 50 µM was used to confirm NMDAR dependence of plateau potentials. SK channel positive allosteric modulator NS309 (10 µM, *Tocris*) was included in a subset of experiments to restore appropriate duration of dendritic integration in Y647S$^{+/-}$ mice. SK2 channel blocker Leidab7 (100 nM) was included in a subset of experiments along with NS309 to determine the contribution of SK2 channels to NS309's effect.

3. *<u>Cellular NMDA currents</u>*

   To examine the impact on neuron-wide NMDARs, whole-cell currents evoked by bath application of NMDA (20 µM, *Tocris*) for 30s were measured in voltage clamp at -75 mV, using K-gluconate patch solution and a modified ACSF to reduce NMDAR $Mg^{2+}$ blockade (0.5 mM $MgSO_4$ & 5 mM KCl) in the presence of AMPAR blockers (CNQX 20 µM). NMDAR antagonist d-APV (50 µM) was used in a subset of all experiments to verify NMDAR mediation of whole-cell currents.

4. *<u>NMDAR EPSC IV curves</u>*

   K- gluconate patch pipettes with 5 mM QX-314 were used to measure NMDAR EPSCs at positive membrane potentials while also keeping other ion channel contributions intact. Mainly this allowed us to assess SK channel mediated negative feedback of NMDARs. I-V curves were generated by recording NMDAR-EPSCs in response to basal dendritic electrical stimulation in the presence of AMPAR and GABAR blockers at different membrane potentials (-85 to -5 mV).

**Wide-field calcium imaging analysis**

Image acquisition was binned every 2 pixels, and the resulting 1024 x 1024-pixel image was subdivided into square regions of 100 x 100 pixels (~220 x 220 $\mu m^2$). Fluorescence intensity change (ΔF/F) over baseline in each subregion of the slice was quantified, and kinetics and spatial spread of population activity were measured. ImageJ and Matlab were used for analysis. Half width (ms) of calcium signals were determined at the region with the highest ΔF/F response. The area under the ΔF/F curve (AUC) for 2s following the electrical stimulus was determined for all regions in the slice and the total number of regions with AUC > 1 Standard Deviation from the mean was taken as the total activated area of the slice. GABAergic blockers (Picrotoxin 20 µM, CGP52432 1 µM) were used to mimic seizure-promoting conditions. To examine persistent activity, two single pulse stimuli were delivered 1.5 seconds apart ~4 seconds following the initial 10 pulse stimulation. Epileptiform activity in Y647S$^{+/-}$ mice was characterized by a large amplitude response to the single pulse stimulus that persisted continuously with recurrent excitation even in the absence of any stimulation in the intervening period.

### Supplemental Results

#### Electrophysiological properties

Intrinsic electrophysiological properties of neurons including resting membrane potential, input resistance, minimum current required to elicit an action potential (rheobase), and action potential threshold were identical in WT and *Grin1* Y647S$^{+/-}$ mice (**Supplementary Table S1**). Membrane capacitance showed a small decrease in Y647S$^{+/-}$ neurons. Input-output curve characterizing the average action potential frequency at increasing amplitude of current injection revealed a small increase in overall excitability of Y647S$^{+/-}$ neurons (19% increase in slope of input-output curve, $P < 0.01$, **Supplementary Fig S1**). Somatic afterhyperpolarization and sag ratio were not different in Y647S$^{+/-}$ neurons (data not shown).

#### Magnesium Threonate improves weight gain and reduces hyperactivity in *Grin1* Y647S$^{+/-}$ mice

Y647S$^{+/-}$ mice are known to be underweight and hyperactive[4] and we examined whether Magnesium-L- Threonate (MgT) treatment improved these phenotypes in addition to reducing seizure occurrence (**Supplementary Fig S2**). Percentage weight gain in Y647S$^{+/-}$ mice at 7 weeks of treatment showed an improvement (67% of WT, $t_{(11)}=2.77$, $P = 0.02$) compared to control Y647S$^{+/-}$ mice (33% of WT). Total distance traveled on average was similar across WT, Y647S$^{+/-}$, and Y647S$^{+/-}$ + MgT mice during open field behavioural assessment but more variable in Y647S$^{+/-}$ mice. Running velocity was higher in Y647S$^{+/-}$ mice compared to WT (P $<10^{-4}$) and improved in MgT-treated Y647S$^{+/-}$ mice ($P <10^{-4}$), showing no significant difference from WT ($P = 0.47$, Kruskal Wallis test). MgT treatment appears beneficial for increasing body weight and reducing hyperactivity in Y647S$^{+/-}$ mice.

### Supplemental Table S1

| | WT (n = 16) | Y647S$^{+/-}$ (n = 19) | T-test statistics |
|---|---|---|---|
| Resting membrane potential (mV) | -83 ± 2 | -84 ± 1 | $t_{(33)} = 0.52$, $P = 0.61$ |
| Input resistance (MΩ) | 124 ± 4 | 119 ± 9 | $t_{(33)} = 0.36$, $P = 0.76$ |
| Membrane Capacitance (pF) | 112 ± 4 | 95 ± 6 | $t_{(33)} = 2.27$, $*P = 0.03$ |
| Action potential threshold (mV) | -48 ± 1 | -50 ± 1 | $t_{(33)} = 1.23$, $P = 0.23$ |
| Rheobase (pA) | 122 ± 13 | 110 ± 10 | $t_{(31)} = 0.73$, $P = 0.47$ |

**Supplementary Table 1. Intrinsic electrophysiological properties of layer 5 pyramidal neurons.** Layer 5 pyramidal neurons in the medial prefrontal cortex of WT and *Grin1* Y647S$^{+/-}$ mice show similar intrinsic electrophysiological properties, except for membrane capacitance, which is significantly decreased in *Grin1* Y647S$^{+/-}$ neurons.

### Supplemental Figure S1

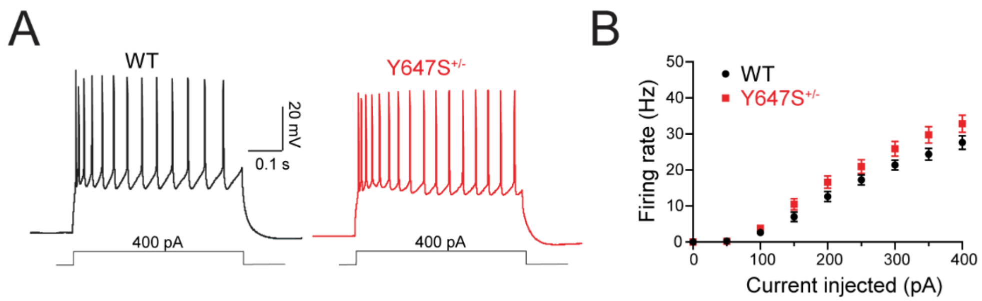


**Intrinsic excitability is slightly increased in *Grin1* Y647S$^{+/-}$ neurons. A,** Action potentials in layer 5 pyramidal neurons with 400 pA constant current injection for 0.5 s in WT and *Grin1* Y647S$^{+/-}$. **B,** Input output curve showing action potential frequency at different current injections. Difference in slope of input-output curve: $F_{(1, 311)} = 6.68$, $P = 0.009$. There is no significant difference in firing rate at any individual current injection (Sidak's post hoc after 2-way ANOVA).

**Supplemental Figure S2**

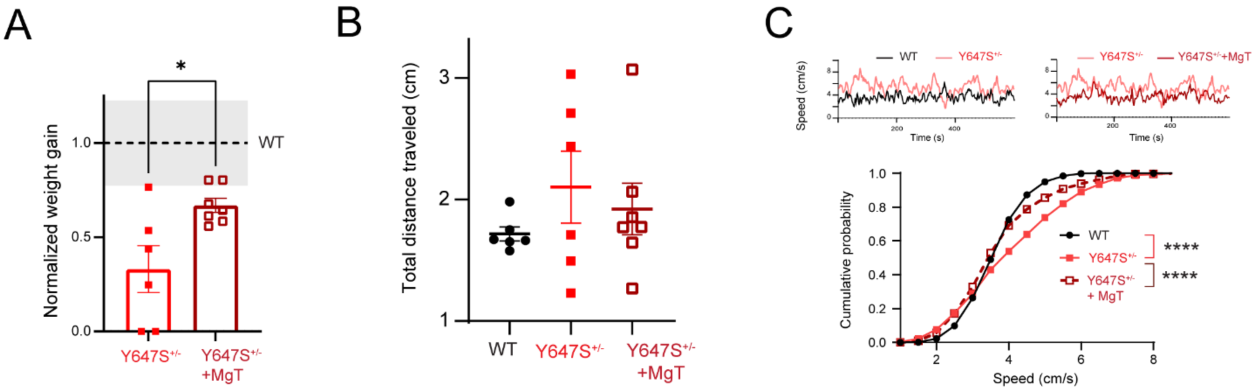


**Magnesium threonate improves weight gain and restores normal running velocity in *Grin1* Y647S$^{+/-}$ mice at 7 weeks of treatment. A,** Normalized weight gain is significantly reduced in Y647S$^{+/-}$ compared to WT ($t_{(10)} = 2.45$, $P = 0.034$), but is improved in MgT treated Y647S$^{+/-}$ mice compared to untreated controls ($t_{(11)} = 2.77$, $P = 0.018$, *$P < 0.05$). Standard error in WT weight gain is shown as the shaded grey rectangle. **B,** Total distance traveled in 10 minutes is not significantly different across WT, Y647S$^{+/-}$, and Y647S$^{+/-}$ +MgT (One-way ANOVA: F (2, 16) = 0.77, P = 0.48) but is more variable in Y647S$^{+/-}$ mice (Brown-Forsythe test: $F_{(2, 16)} = 4.23$, $P = 0.034$). **C,** Velocity profile comparing example WT and Y647S$^{+/-}$ mice (Top left), and the same Y647S$^{+/-}$ with an MgT-treated Y647S$^{+/-}$ mouse (Top right). Cumulative distribution of velocity in WT, Y647S$^{+/-}$, and Y647S$^{+/-}$+MgT mice (WT vs Y647S$^{+/-}$: $P < 10^{-4}$, Y647S$^{+/-}$ vs Y647S$^{+/-}$ + MgT: $P < 10^{-4}$, WT vs Y647S$^{+/-}$ + MgT: $P = 0.47$, Dunn's post hoc). ****$P < 10^{-4}$.
